# Identification of Candidate COVID-19 Therapeutics using hPSC-derived Lung Organoids

**DOI:** 10.1101/2020.05.05.079095

**Authors:** Yuling Han, Liuliu Yang, Xiaohua Duan, Fuyu Duan, Benjamin E. Nilsson-Payant, Tomer M. Yaron, Pengfei Wang, Xuming Tang, Tuo Zhang, Zeping Zhao, Yaron Bram, David Redmond, Sean Houghton, Duc Nguyen, Dong Xu, Xing Wang, Skyler Uhl, Yaoxing Huang, Jared L. Johnson, Jenny Xiang, Hui Wang, Fong Cheng Pan, Lewis C. Cantley, Benjamin R. tenOever, David D. Ho, Todd Evans, Robert E. Schwartz, Huanhuan Joyce Chen, Shuibing Chen

## Abstract

The SARS-CoV-2 virus has caused already over 3.5 million COVID-19 cases and 250,000 deaths globally. There is an urgent need to create novel models to study SARS-CoV-2 using human disease-relevant cells to understand key features of virus biology and facilitate drug screening. As primary SARS-CoV-2 infection is respiratory-based, we developed a lung organoid model using human pluripotent stem cells (hPSCs) that could be adapted for drug screens. The lung organoids, particularly aveolar type II cells, express ACE2 and are permissive to SARS-CoV-2 infection. Transcriptomic analysis following SARS-CoV-2 infection revealed a robust induction of chemokines and cytokines with little type I/III interferon signaling, similar to that observed amongst human COVID-19 pulmonary infections. We performed a high throughput screen using hPSC-derived lung organoids and identified FDA-approved drug candidates, including imatinib and mycophenolic acid, as inhibitors of SARS-CoV-2 entry. Pre- or post-treatment with these drugs at physiologically relevant levels decreased SARS-CoV-2 infection of hPSC-derived lung organoids. Together, these data demonstrate that hPSC-derived lung cells infected by SARS-CoV-2 can model human COVID-19 disease and provide a valuable resource to screen for FDA-approved drugs that might be repurposed and should be considered for COVID-19 clinical trials.

---

The ongoing COVID-19 pandemic is an unprecedented global event that requires the immediate deployment of effective clinical therapeutics while vaccine candidates are identified and tested. Arguably the most rapid means of addressing this issue is through the repurposing of existing drugs that are FDA-approved which may indirectly interfere with aspects of SARS-CoV-2 biology. While this strategy is being pursued, most high throughput screens focus on the use of transformed cell lines which fail to capture the physiologically relevant dynamics of a SARS-CoV-2 infection. In an effort to improve on these cell lines, we developed and describe here lung organoids as an improved in vitro platform for screening purposes.

In the last several years, a series of protocols have been reported to direct hPSC differentiation to various lung lineages^1–14^. We differentiated hPSCs to lung organoids using a previously reported stepwise strategy^4,15^, including the progressive differentiation first into definitive endoderm (DE), followed by specification to anterior foregut endoderm (AFE), AFE/lung progenitor cells (LPs), and finally lung organoids (**Extended Data Fig. 1)**. Single cell transcriptomic profiles were generated and analyzed in the differentiated lung organoids at day 50 and identified alveolar type II (AT2) cells (SP-B^+^, SP-D^+^, ABCA3^+^), alveolar type I (AT1) cells (PDPN^+^APQ5^+^), stromal cells, and proliferating cells (**Fig. 1a, 1b** and **Extended Data Fig. 2b**). We also detected a low number of pulmonary neuroendocrine cells (ASCL1^+^, CALCA^+^) and airway epithelial cells (**Fig. 1a** and **Extended Data Fig. 2a, 2b**). ACE2, the putative receptor for SARS-CoV-2^16^, is mainly detected in cluster 1, which represents AT2 cells (**Fig. 1c, 1d**). TMPRSS2, a key transmembrane protease for SARS-CoV-2 infection^16^, is also enriched in AT2 cells and AT1 cells (**Fig. 1c, 1d**). Consistent with scRNA-seq data of adult lung^17^, ACE2 and TMPRSS2 expression are detected in only a subset of the AT2 cells, likely due to the depth limitation of 10X scRNA-seq. Immunostaining results further validated that ACE2 is expressed in SP-B^+^/SP-C^+^ AT2 cells (**Fig. 1e**).

**Figure 1.**
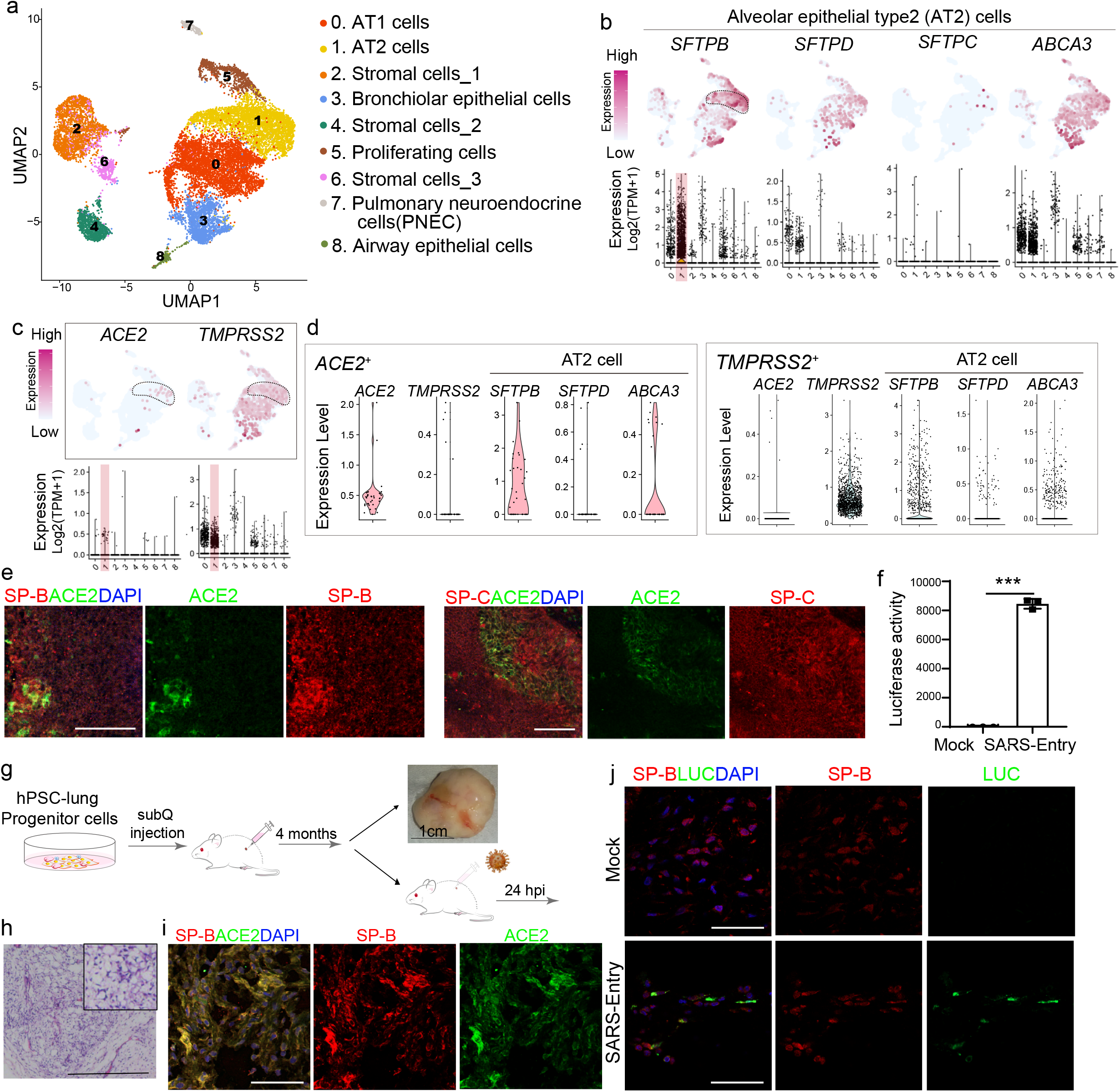
hPSC-derived lung organoids express ACE2 and are permissive to SARS-CoV-2 pseudo-entry virus infection both *in vitro* and *in vivo*. **a.** UMAP of hPSC-derived lung organoids, which contain 14,263 hPSC-derived lung epithelial cells (EPCAM^+^, UMI count>0), colored and annotated with clusters 0-8. AT2 cells, Alveolar Epithelial Type 2 cells. AT1 cells, Alveolar Epithelial Type 1 cells. **b.** Putative AT2 markers in each cluster in UMAPs. Relative expression level of each marker gene range from low (light blue) to high (pink) as indicated. Individual cells positive for lung cell markers are denoted by red dots. The violin plot shows the expression level (log2(TPM+1)) of each indicated gene in each cluster. **c.** UMAP of *ACE2* and *TMPRSS2* expression in AT2 cells. **d.** Violin plots of *ACE2* and *TMPRSS2* expression in cells expressing AT2 markers including *SFTPB, SFPTD* and *ABCA3*. **e.** Immunostaining of hPSC-derived lung organoids detected the co-expression of ACE2 in SP-B^+^ or SP-C^+^ AT2-like cells. Scale bars= 25 μm. **f.** Luciferase activity of hPSC-derived lung organoids either mock-infected or infected with SARS-CoV-2 pseudo-entry virus at 24 hpi (MOI=0.01). **g.** Schematic of the experimental flowchart for the mouse xenograft model formed with hPSC-derived lung cells. Briefly, the day 25 lung progenitor cells were injected subcutaneously in NSG mice and xenografts were collected after 4 months of transplantation. subQ injection, subcutaneous injection. The cells within the xenografts were isolated and analyzed by immunostaining and infection of SARS-CoV-2 pseudoentry virus. **h,** Representative image of Hematoxylin and Eosin staining on the xenograft lung tissue that shows the typical alveolar region. Scale Bars= 200μm. **i,** Immunostaining of hPSC-derived lung xenografts detected the expression of ACE2 in SP-B^+^ cells. Scale bars= 25 μm. **j,** Immunostaining of hPSC-derived lung xenografts at 24 hpi (1X10^4^ FFU) detected the coexpression of luciferase (LUC) in SP-B^+^ cells. Scale bars= 25 μm. Data was presented as mean ±STDEV. *P* values were calculated by unpaired two-tailed Student’s t test. \**P* < 0.05, \*\**P* < 0.01, and \*\*\**P* < 0.001.

To determine the relative permissiveness of hPSC-derived lung organoids to SARS-CoV-2 viral entry, we first used a vesicular stomatitis virus (VSV) based SARS-CoV-2 pseudo-entry virus, for which the backbone was provided by a VSV-G pseudo-typed ΔG-luciferase virus with the SARS-CoV-2 Spike protein incorporated at the surface of the viral particle (See Methods for details)^18,19^. Robust luciferase activity was readily detected in the infected hPSC-derived lung organoids (**Fig. 1f**).

To generate an *in vivo* model using human lung organoids, we implanted subcutaneously day 25 lung progenitor cells in immuno-deficient NSG mice (**Fig. 1g**). Within 4 months the xenografts developed organized alveolar-like structures (**Fig. 1h**). Immunostaining confirms the existence of SP-B^+^ AT2 cells, which co-express ACE2 (**Fig. 1i**). Infection of SARS-CoV-2 pseudo-entry virus was tested in this mouse model carrying hPSC-derived lung xenografts. Expression of luciferase from the SARS-CoV-2 pseudo-entry virus was detected by immunofluorescence staining 24 hours after intra-xenograft inoculation (1X10^4^ FFU). LUC is mainly detected in SP-B^+^ AT2 cells (**Fig. 1j**).

The potential of the hPSC-derived lung organoid platform to model COVID-19 was then tested by infection with SARS-CoV-2 virus in the cultures. 24 hours post inoculation (hpi) with the SARS-CoV-2 virus (USA-WA1/2020, MOI=0.01), qRT-PCR using primers targeting N sgRNA transcripts confirmed that a significant amount of replicating viral RNA was detected in the infected lung organoids (**Fig. 2a**). Immunostaining confirmed the detection of SARS-S protein inthe infected lung organoids (**Fig. 2b**). At 24 hpi, RNA-seq was performed on mock and SARS-CoV-2 infected cells. Alignment with the viral genome confirmed robust viral replication in hPSC-derived lung organoids (**Fig. 2c**). Moreover, plotting these datasets by principle component analysis (PCA) suggested that the infected lung organoids clustered distinctly compared to mock-infected lung organoids (**Fig. 2d**). Volcano plots of SARS-CoV-2 infected hPSC-derived lung organoids compared to mock treatment revealed robust induction of chemokines and cytokines with no detectable levels of type I and III IFNs (**Fig. 2e**). Gene set enrichment analysis (GSEA) comparing mock-infected versus SARS-CoV-2 infected lung organoids revealed over-represented pathway networks including TNF signaling, IL-17 signaling, chemokine signaling pathway, and cytokine-cytokine receptor interaction (**Fig. 2f**). These profiles were further compared with primary tissues from healthy and COVID-19 patients. Compared with healthy lung, lung tissues of COVID-19 patients revealed robust induction of chemokines, including *CXCl2, CCL2, CXCL3* as well as *IL1A, BCRC3, AADAC*, and *ATPB4* (**Fig. 2g**), which is markedly similar to SARS-CoV-2 infected lung organoids. Finally, IL-17 signaling was also found to be significantly changed in lung tissues of COVID-19 patients, which is consistent with SARS-CoV-2 infected lung organoids (**Fig. 2h**).

**Figure 2.**
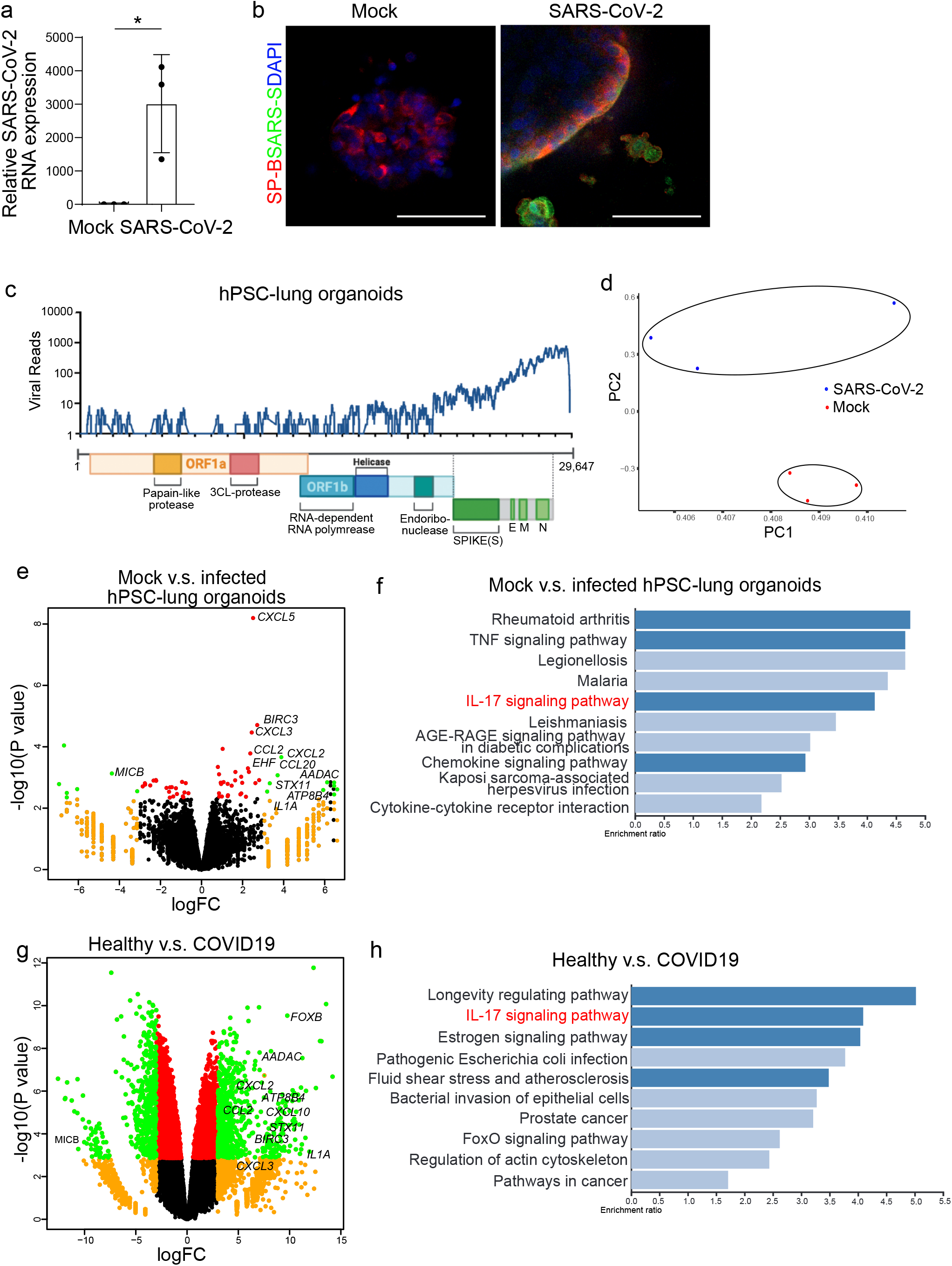
Transcriptomic analysis of SARS-CoV-2 infected hPSC-derived lung organoids demonstrates robust SARS-CoV-2 replication, upregulation of chemokine expression with no upregulation of Type I/III IFN signaling. **a,** Relative SARS-CoV-2 R viral NA expression in hPSC-derived lung organoids. Total viral RNA from infected hPSC-derived lung organoids (MOI=0.01) was analyzed by qRT-PCR for the presence of N sgRNA transcripts relative to ACTB. **b,** Immunostaining of hPSC-derived lung organoids at 24 hpi (SARS-CoV-2, MOI=0.01) detected the expression of SARS-S in SP-B^+^ cells. Scale bars= 25 μm. **c,** Alignment of the transcriptome with the viral genome in SARS-CoV-2 infected hPSC-derived lung organoids. Schematic below shows the SARS-CoV-2 genome. **d,** PCA plot of mock-infected or SARS-CoV-2 infected hPSC-derived lung organoids. **e,** Volcano plot analysis of differential expression of SARS-CoV-2 infected hPSC-derived lung organoids versus mock infection. Individual genes are denoted by gene name. **f,** Gene over-representation analysis on KEGG pathway database of SARS-CoV-2 infected hPSC-derived lung organoids versus mock infection. **g,** Volcano plot analysis of differential expression of lung biopsy from COVID-19 versus healthy patients. Individual genes are denoted by gene name. **h.** Gene over-representation analysis on KEGG pathway database of lung biopsy from COVID-19 versus healthy patients (GSE147507)^23^. Data was presented as mean ± STDEV. *P* values were calculated by unpaired two-tailed Student’s t test. \**P* < 0.05, \*\**P* < 0.01, and \*\*\**P* < 0.001.

To identify drug candidates capable of blocking SARS-CoV-2 pseudo-virus infection, hPSC-derived lung organoids were deposited onto 384-well plates. After six hour of incubation, organoids were treated at 10 μM with a library of FDA-approved drugs (the Prestwick collection). Two hour post-treatment, the organoids were innoculated with SARS-CoV-2 pseudo-entry virus at MOI=0.01. At 24 hpi, the organoids were analyzed for luciferase activity. The wells in which Z score<-2 were chosen as primary hit drugs (**Fig. 3a**). The hits were evaluated for efficacy and cytotoxicity at different concentrations. Four drugs were confirmed to block luciferase activity in a dose-dependent manner, independent of cytotoxicity, including imatinib (EC50=4.86 μM, IC50=37.3 μM **Fig. 3b, 3e**), mycophenolic acid (MPA, EC50=0.15 μM, **Fig. 3c, 3f**), quinacrine dihydrochloride (QNHC, EC50=2.83 μM, IC50=22 μM, **Fig. 3d, 3g**), chloroquine (EC50=3.85 μM), and prochlorperazine (EC50=23.7 μM, IC50=30 μM) (**Extended Data Fig. 3**). Interestingly, three hit compounds, MPA, QNHC, and chloroquine, were also identified from our independent screen using hPSC-derived colonic organoids^20^. Immunostaining confirmed a significant diminishment of LUC^+^ cells detected among SP-C^+^ AT2 cells in lung organoids treated with 10 μM imatinib, 3 μM MPA or 4.5 μM QNHC at 24 hpi (**Fig. 3h**).

**Figure 3.**
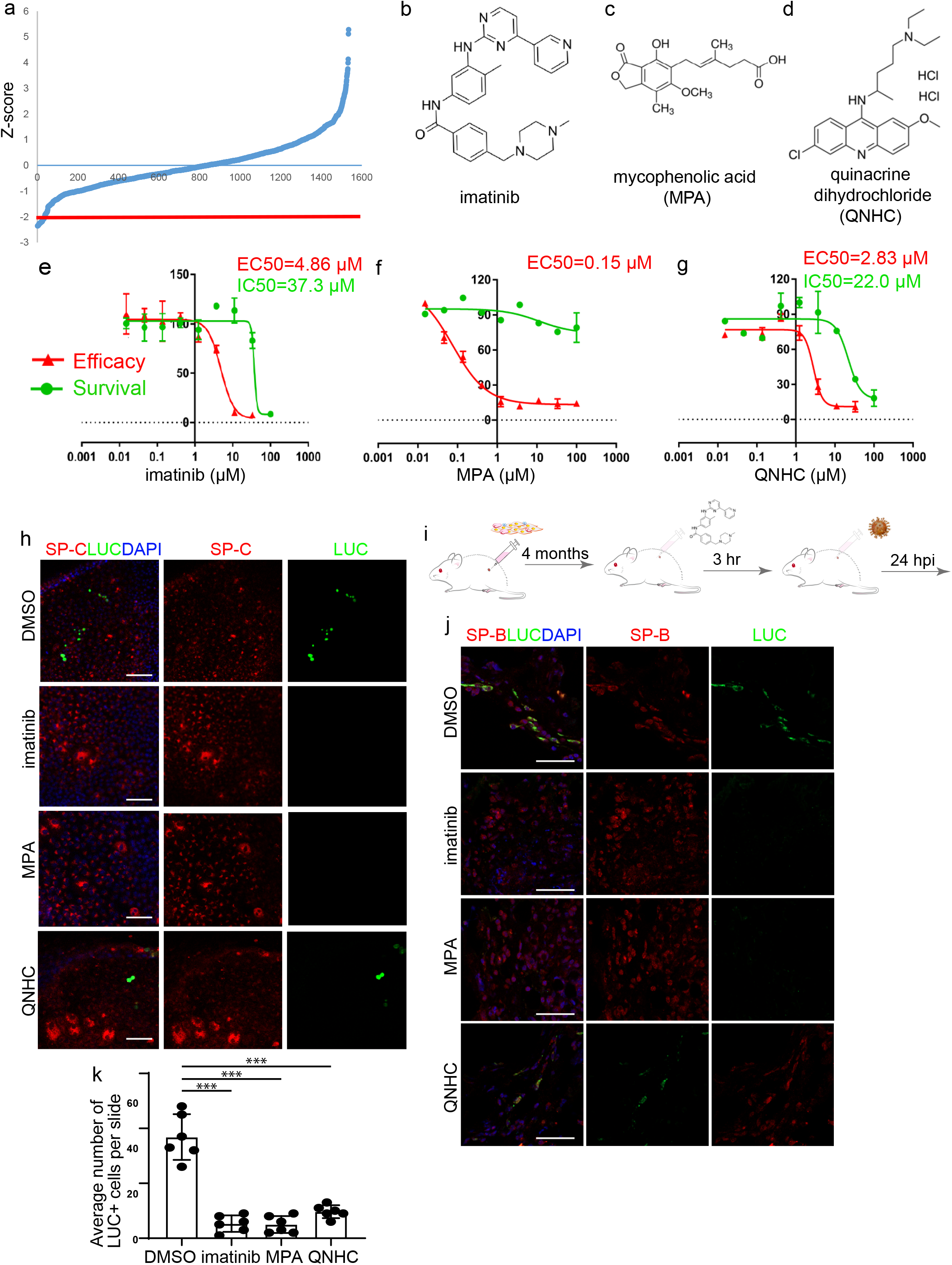
A hPSC-derived lung organoid-based high throughput chemical screen identifies three FDA-approved drug candidates that block SARS-CoV-2 entry. **a,** Primary screening results. **b-d,** Chemical structure of imatinib (b), mycophenolic acid (MPA, c), and quinacrine dihydrochloride (QNHC, d). **e-g,** Efficacy and toxicity curves of imatinib (e), MPA (f), and QNHC (g). Data is presented as mean ± STDEV. N=3. **h,** Immunostaining of LUC^+^ cells in imatinib, MPA, and QNHC-treated hPSC-derived lung organoids at 24 hpi (MOI=0.01). Scale bar = 25 μm. **i,** Scheme of *in vivo* drug treatment. **j-k,** Immunostaining (j) and quantification (k) of hPSC-derived lung xenografts of mice treated with 400 mg/kg imatinib, 50 mg/kg MPA, and 25 mg/kg QNHC at 24 hpi (1X10^4^ FFU). Scale bars= 25 μm. N= 6 xenografts per condition 5 slides per xenograft. Data was presented as mean ± STDEV. *P* values were calculated by unpaired two-tailed Student’s t test. \**P* < 0.05, \*\**P* < 0.01, and \*\*\**P* < 0.001.

To evaluate the drug activities *in vivo*, we used humanized mice carrying hPSC-derived lung xenografts after 4 months maturation *in vivo* (**Fig. 3i**). The humanized mice were treated with 400 mg/kg imatinib mesylate, 50 mg/kg MPA, or 25 mg/kg QNHC. 3 hours post-treatment, SARS-CoV-2 pseudo-entry virus (1X10^4^ FFU) was delivered by intra-xenograft inoculation. At 24 hpi, luciferase staining was detected in mice treated with vehicle. The number of LUC^+^ cells was significantly decreased in mice treated with imatinib mesylate, MPA or QNHC (**Fig. 3j, 3k**).

Finally, hPSC-derived lung organoids pre-treated with 10 μM imatinib, 3 μM MPA or 4.5 μM QNHC were infected with SARS-CoV-2 virus at MOI=0.5. At 24 hpi, qRT-PCR confirmed significantly decreased replicating viral RNA in lung organoids treated with imatinib, MPA or QNHC (**Fig. 4a**). Immunostaining confirmed a significant loss of SARS-CoV-2^+^ cells in imatinib, MPA or QNHC-treated hPSC derived-lung organoids (**Fig. 4b**). To determine the therapeutic potential of imatinib, MPA and QNHC, hPSC-derived lung organoids were infected with SARS-CoV-2 virus (MOI=0.5). Three hours later, lung organoids were treated with 10 μM imatinib, 3 μM MPA or 4.5 μM QNHC. At 24 hpi, both viral RNA (**Fig. 4c**) and SARS-CoV-2^+^ cells (**Fig. 4d**) were significantly decreased in the lung organoids treated with each drug, highlighting the therapeutic potential. Transcriptional profiling was applied to compare DMSO and imatinib-treated lung organoids, and PCA plots showed these clustered separately (**Fig. 4e**). Volcano plots and GSEA analysis highlight the change of pathways caused by imatinib, related to fatty acid biosynthesis, steroid biosynthesis, fatty acid metabolism, and PPAR signaling pathway (**Fig. 4f, 4g**). Viruses have been known to target lipid signaling, synthesis, and metabolism to remodel their host cells into an optimal environment for their replication. Fatty acid is involved in multiple steps of viral circle, including membrane fusion during the entry process, virion envelopment during particle maturation, as well as virus replication^21^. The fact that fatty acid biosynthesis and metabolism pathways are changed in the imatinib-treated lung organoids suggests that imatinib might also affect virus replication and particle maturation. Finally, qRT-PCR and western blotting experiments confirmed the ability of imatinib to block anti-SARS-CoV-2 activity in Vero cells (**Extended Data Fig. 4**).

**Figure 4.**
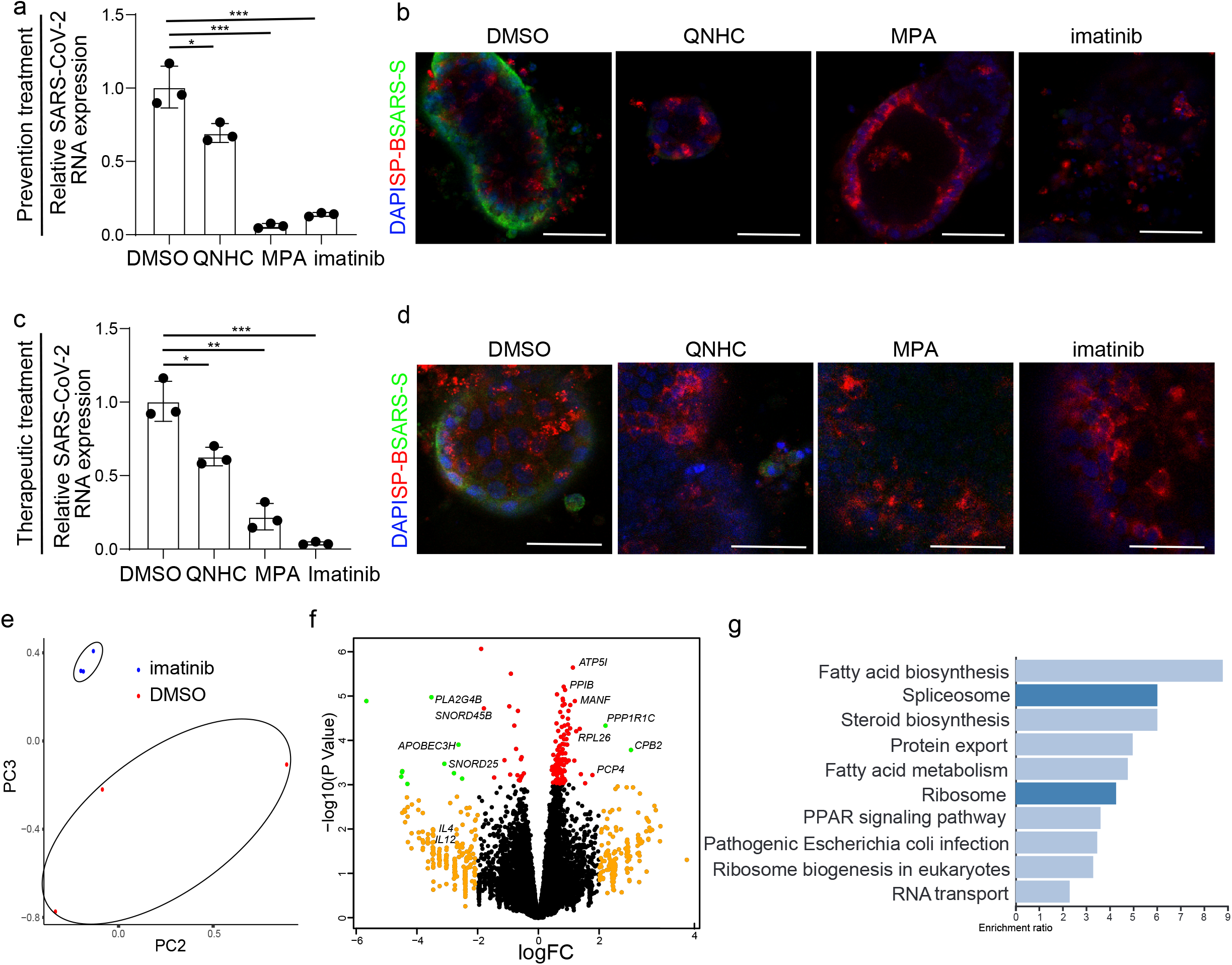
Imatinib, mycophenolic acid, and quinacrine dihydrochloride each block the entry and spreading of SARS-CoV-2 virus. **a,** Relative SARS-CoV-2 viral RNA expression in hPSC-derived lung organoids pre-treated with 10 μM imatinib, 3 μM MPA or 4.5 μM QNHC at 24 hpi of SARS-CoV-2 virus (MOI=0.5). Total viral RNA from infected hPSC-derived lung organoids (MOI=0.01) was analyzed by qRT-PCR for the presence of N sgRNA transcripts relative to ACTB. **b,** Immunostaining of SARS-CoV-2 Spike protein (SARS-S) and SP-B in imatinib, MPA, or QNHC treated hPSC-derived lung organoids at 24 hpi (MOI=0.5). Scale bars = 25 μm. **c,** Relative SARS-CoV-2 viral RNA expression at 24 hpi of hPSC-derived lung organoids infected with SARS-CoV-2 virus (MOI=0.5) and three hours later followed by 10 μM imatinib, 3 μM MPA or 4.5 μM QNHC treatment. Total RNA from infected hPSC-derived lung organoids (MOI=0.5) was analyzed by qRT-PCR for the presence of N sgRNA transcripts relative to ACTB. **d,** Immunostaining of SARS-S and SP-B at 24 hpi of hPSC-derived lung organoids infected with SARS-CoV-2 virus (MOI=0.5) and three hours later followed by 10 μM imatinib, 3 μM MPA or 4.5 μM QNHC treatment. Scale bars = 100 μm. **e,** PCA plot of hPSC-derived lung organoids pretreated with DMSO or 10 μM imatinib at 24 hpi of SARS-CoV-2 virus. **f,** Volcano plot analysis of differential expression of hPSC-derived lung organoids pretreated with DMSO or 10 μM imatinib at 24 hpi of SARS-CoV-2 virus. Individual genes are denoted by gene name. **g,** Gene over-representation analysis on KEGG pathway database of differential expression of hPSC-derived lung organoids pretreated with DMSO or 10 μM imatinib at 24 hpi of SARS-CoV-2 virus. Data was presented as mean ± STDEV. *P* values were calculated by unpaired two-tailed Student’s t test. \**P* < 0.05, \*\**P* < 0.01, and \*\*\**P* < 0.001.

The lung is the most vulnerable target organ for the SARS-CoV-2 virus, and respiratory failure is the primary disease outcome for COVID-19. Yet the primary model currently used for SARS-CoV-2 studies are African green monkey kidney derived Vero cells, which have clear limitations for modeling complex human pulmonary or other organ systems. Therefore, the development of physiologically relevant human cell models to study SARS-CoV-2 infection is critically important. Here, we present an hPSC-derived lung organoid platform, including SP-B^+^ AT2 cells that express ACE2 and TMPRSS2, two key factors involved in SARS-CoV-2 infection, which is consistent with the previous reports^22^. RNA-seq of infected organoids revealed upregulation of cytokine/chemokine signaling, which phenocopies the cytokine and chemokine changes observed in primary human COVID-19 pulmonary infection^23^. Finally, we used the hPSC-derived lung organoids in a high throughput screen for FDA-approved drugs. We identified several drugs that decreased the luciferase activity of SARS-CoV-2 pseudo-entry virus including imatinib, MPA and QNHC, both *in vitro* and *in vivo*. The anti-viral activity of these drugs was further validated against patient-derived SARS-CoV-2 virus.

Imatinib is an inhibitor of a number of tyrosine kinase enzymes, including Abl, c-kit, PDGF-R and others. Previous studies have also suggested imatinib as a potent inhibitor of SARS and MERS coronavirus fusion proteins^24^. Using HIV SARS-S and MERS-S pseudotyped virions, imatinib was shown to substantially block coronavirus S protein-induced fusion and prevent endosomal entry^25^. Imatinib has been widely used to treat chronic myelogenous leukemia and other cancers, and should be considered for repurposing as a drug candidate for COVID-19 patients. Very recently, three clinical trials (ClinicalTrials.gov Identifier: NCT04346147, NCT04357613, NCT04356495) were registered to apply imatinib to treat COVID-19 patients. Our study provides experimental data to support these trials.

## Methods

### hPSC lung differentiation

Protocols for maintenance of hPSCs and generation of lung cells were slightly modified from previous studies^4,15^. The hESC line RUES2 was cultured on irradiated mouse embryonic fibroblasts (Global Stem, cat. no. GSC-6001G) at a density of 20,000-25,000 cells/cm^2^ in a medium of DMEM/F12, 20% knockout serum replacement (Life Technologies), 0.1 mM β-mercaptoethanol (Sigma Aldrich) and 20 ng/ml bFGF (R&D Systems), and medium was changed daily. hESC cultures were maintained in an undifferentiated state at 37 °C in a 5% CO2/air environment until stem cells reached about 90% confluence.

hESC differentiation into endoderm was performed in serum-free differentiation (SFD) medium of DMEM/F12 (3:1) (Life Technologies) supplemented with N2 (Life Technologies), B27, 50 μg/ml ascorbic acid, 2 mM Glutamax, 0.4 μM monothioglycerol, 0.05% BSA at 37 °C in a 5% CO2/5% O2/95% N2 environment. hESCs were treated with Accutase and plated onto low attachment 6-well plates (Corning Incorporated, Tewksbury MA), resuspended in endoderm induction medium containing 10 μM Y-27632, 0.5 ng/ml human BMP-4, 2.5 ng/ml human bFGF, 100 ng/ml human Activin A, for 72-84 hours dependent on the formation rates of endoderm cells. On day 3 or 3.5, the endoderm bodies were dissociated into single cells using 0.05% Trypsin/0.02% EDTA and plated onto fibronectin-coated, 24-well tissue culture plates (~100,000–150,000 cells/well). For induction of anterior foregut endoderm, the endoderm cells were cultured in SFD medium supplemented with 1.5 μM dorsomorphin dihydrochloride (R&D Systems) and 10 μM SB431542 (R&D Systems) for 36-48 h, and then switched to 36-48 h of 10 μM SB431542 and 1 μM IWP2 (R&D Systems) treatment. For induction of early stage lung progenitor cells (day 6–15), the resulting anterior foregut endoderm was treated with 3 μM CHIR99021, 10 ng/ml human FGF10, 10 ng/ml human FGF-7, 10 ng/ml human BMP-4 and 50-60 nM all-trans retinoic acid (ATRA), in SFD medium for 8–10 d. The day 10–15 cultures were maintained in a 5% CO2/air environment. On days 15 and 16, the lung field progenitor cells were replated after one minute trypsinization onto fibronectin-coated plates, in the presence of SFD containing either a combination of five factors (3 μM CHIR99021, 10 ng/ml human FGF10, 10 ng/ml human FGF7, 10 ng/ml human BMP-4, and 50 nM ATRA), or three factors (3 μM CHIR99021, 10 ng/ml human FGF10, 10 ng/ml human FGF7) for day 14-16. Day 16–25 cultures of late stage lung progenitor cells were maintained in SFD media containing 3 μM CHIR99021, 10 ng/ml human FGF10, 10 ng/ml human FGF7, in a 5% CO2/air environment. For differentiation of mature lung cells (day 25 to 55), cultures were re-plated after brief trypsinization onto 3.3% Matrigel-coated 24-well plates in SFD media containing maturation components containing 3 μM CHIR99021, 10 ng/ml human FGF-10; 10 ng/ml human FGF7, and DCI (50 nM Dexamethasone, 0.1 mM 8-bromo-cAMP (Sigma Aldrich) and 0.1 mM IBMX (3,7-dihydro-1-methyl-3-(2-methylpropyl)-1H-purine-2,6-dione) (Sigma Aldrich)). 1 μM DAPT was added to the maturation media for induction of pulmonary neuroendocrine cells (PNECs) and Tuft cells. The protocol details are summarized in Figure S1A.

### Cell Lines

HEK293T (human [*Homo sapiens*] fetal kidney) and Vero E6 (African green monkey [*Chlorocebus aethiops*] kidney) were obtained from ATCC (https://www.atcc.org/). Cells were cultured in Dulbecco’s Modified Eagle Medium (DMEM) supplemented with 10% FBS and 100 I.U./mL penicillin and 100 μg/mL streptomycin. All cell lines were incubated at 37°C with 5% CO2.

### SARS-CoV-2-Pseudo-Entry Viruses

Recombinant Indiana VSV (rVSV) expressing SARS-CoV-2 spikes was generated as previously described ^18,19,26^. HEK293T cells were grown to 80% confluency before transfection with pCMV3-SARS-CoV2-spike (kindly provided by Dr. Peihui Wang, Shandong University, China) using FuGENE 6 (Promega). Cells were cultured overnight at 37°C with 5% CO2. The next day, the media was removed and VSV-G pseudotyped ΔG-luciferase (G*ΔG-luciferase, Kerafast) was used to infect the cells in DMEM at an MOI of 3 for 1 hr before washing the cells with 1X DPBS three times. DMEM supplemented with 2% FBS and 100 I.U. /mL penicillin and 100 μg/mL streptomycin was added to the infected cells and they were cultured overnight as described above. The next day, the supernatant was harvested and clarified by centrifugation at 300xg for 10 min before aliquoting and storing at −80°C.

### SARS-CoV-2 Viruses

SARS-CoV-2, isolate USA-WA1/2020 (NR-52281) was deposited by the Center for Disease Control and Prevention and obtained through BEI Resources, NIAID, NIH. SARS-CoV-2 was propagated in Vero E6 cells in DMEM supplemented with 2% FBS, 4.5 g/L D-glucose, 4 mM L-glutamine, 10 mM Non-Essential Amino Acids, 1 mM Sodium Pyruvate and 10 mM HEPES as described previously^23^.

All work involving live SARS-CoV-2 was performed in the CDC/USDA-approved BSL-3 facility of the Global Health and Emerging Pathogens Institute at the Icahn School of Medicine at Mount Sinai in accordance with institutional biosafety requirements

### SARS-CoV-2 entry virus infections

For lung organoids, organoids were seeded in 24-well plates, pseudo-typed virus was added for MOI=0.01 and centrifuged the plate at 1200g, 1 hour. At 24 hpi, organoids were fixed for immunohistochemistry or harvested for luciferase assay following the Luciferase Assay System protocol (E1501, Promega)

### SARS-CoV-2 virus infections

hESC-derived lung organoids were infected with SARS-CoV-2 at the indicated MOI and incubated for 24 h at 37°C. Where indicated, hESC-derived lung organoids were pretreated with DMSO, 10 μM imatinib, 3 μM MPA or 4.5 μM QNHC for 3 h prior to infection as well as during the course of infection. Where indicated, hESC-derived lung organoids were treated with DMSO, 10 μM imatinib, 3 μM MPA or 4.5 μM QNHC 3 hpi. At the time point of harvest, cells were washed three times with PBS and harvested for either RNA analysis or immunofluorescence staining.

Approximately 2.5 × 105 Vero E6 cells were pre-treated with DMSO, 10 μM imatinib, 3 μM MPA or 4.5 μM QNHC for 1 h prior to infection with SARS-CoV-2 at an MOI of 0.01 in DMEM supplemented with 2% FBS, 4.5 g/L D-glucose, 4 mM L-glutamine, 10 mM Non-Essential Amino Acids, 1 mM Sodium Pyruvate and 10 mM HEPES. At 24 hpi, cells were washed three times with PBS before harvesting for RNA or protein analysis.

Cells were either lysed in TRIzol for RNA analysis or in RIPA buffer for protein analysis or fixed in 5% formaldehyde for 24 h for immunofluorescent staining, prior to safe removal from the BSL-3 facility.

### Xenograft formation

1 million hESC-derived cells at lung progenitor stage (at day 25) were subcutaneously injected into 6-8 weeks old *NOD.Cg-Prkdcscid Il2rgtm1WjI/SzJ* (NSG) mice (Jackson Laboratory, Bar Harbor, Maine). When xenograft size becomes 1-2 CM^3^, they were sacrificed immediately, necropsy performed, and cells were harvested for further histological or molecular study.

### Immunohistochemistry

Histology on tissues from mice was performed on paraffin-embedded or frozen sections from xenografts and corresponding normal tissues as previously described^27^. Tissues were fixed overnight in 10% buffered formalin and transferred to 70% ethanol, followed by paraffin embedding, or tissues were fixed in 10% buffered formalin and transferred to 30% sucrose, followed by snap frozen in O.C.T (Fisher Scientific, Pittsburgh, PA). Adjacent sections stained with Hematoxylin and Eosin were used for comparison. Living cells in culture were directly fixed in 4% paraformaldehyde for 25 min, followed with 15 min permeabilization in 0.1% Triton X-100. For immunofluorescence, cells or tissue sections were immunostained with primary antibodies at 4°C overnight and secondary antibodies at RT for 1h. The information for primary antibodies and secondary antibodies are provided in Table S3. Nuclei were counterstained by DAPI.

### Western blot

Protein was extracted from cells in Radioimmunoprecipitation assay (RIPA) lysis buffer containing 1X Complete Protease Inhibitor Cocktail (Roche) and 1X Phenylmethylsulfonyl fluoride (Sigma Aldrich) prior to safe removal from the BSL-3 facility. Samples were analysed by SDS-PAGE and transferred onto nitrocellulose membranes. Proteins were detected using rabbit polyclonal anti-GAPDH (Sigma Aldrich, G9545), mouse monoclonal anti-SARS-CoV-2 Nucleocapsid [1C7] and mouse monoclonal anti-SARS-CoV-2 Spike [2B3E5] protein (a kind gift by Dr. T. Moran, Center for Therapeutic Antibody Discovery at the Icahn School of Medicine at Mount Sinai). Primary antibodies were detected using Fluorophore-conjugated secondary goat anti-mouse (IRDye 680RD, 926-68070) and goat anti-rabbit (IRDye 800CW, 926-32211) antibodies. Antibody-mediated fluorescence was detected on a LI-COR Odyssey CLx imaging system and analyzed using Image Studio software (LI-COR).

### qRT-PCR

Total RNA samples were prepared from cells/organoids using TRIzol and Direct-zol RNA Miniprep Plus kit (Zymo Research) according to the manufacturer’s instructions. To quantify viral replication, measured by the expression of sgRNA transcription of the viral N gene, one-step quantitative real-time PCR was performed using SuperScript III Platinum SYBR Green One-Step qRT-PCR Kit (Invitrogen) with primers specific for the TRS-L and TRS-B sites for the N gene as well as ACTB as an internal reference. Quantitative real-time PCR reactions were performed on a LightCycler 480 Instrument II (Roche). Delta-delta-cycle threshold (ΔΔCT) was determined relative to the ACTB and mock infected /treated samples. Error bars indicate the standard deviation of the mean from three biological replicates. The sequences of primers/probes are provided in Table S4.

### Single-cell RNA-seq data analysis

We filtered cells with less than 200 or more than 6000 genes detected as well as cells with mitochondria gene content greater than 30%, and used the remaining 14263 cells for downstream analysis. We normalized the gene expression UMI counts using a deconvolution strategy implemented by the R scran package (v.1.14.1). In particular, we pre-clustered cells using the *quickCluster* function; we computed size factor per cell within each cluster and rescaled the size factors by normalization between clusters using the *computeSumFactors* function; and we normalized the UMI counts per cell by the size factors and took a logarithm transform using the *normalize* function. We identified highly variable genes using the *FindVariableFeatures* function in the R Seurat (v3.1.0) ^28^, and selected the top 3000 variable genes after excluding mitochondria genes, ribosomal genes and dissociation-related genes. The list of dissociation-related genes was originally built on mouse data ^29^; we converted them to human ortholog genes using Ensembl BioMart. We scaled the normalized counts and performed PCA on the highly variable genes using the *ScaleData* and *RunPCA* functions in the R Seurat package ^28^. We selected the top 20 PCs for downstream visualization and clustering analysis. We ran UMAP dimensional reduction using the *RunUMAP* function in the R Seurat package with the number of neighboring points setting to 35 and training epochs setting to 500. We clustered cells into fifteen clusters by constructing a shared nearest neighbor graph and then grouping cells of similar transcriptome profiles using the *FindNeighbors* function and *FindClusters* function (resolution set to 0.2) in the R Seurat package. We identified marker genes for each cluster by performing differential expression analysis between cells inside and outside that cluster using the *FindMarkers* function in the R Seurat package. After reviewing the clusters, we merged them into nine clusters representing nine cell types (AT1 cells, AT2 cells, bronchiolar epithelial cells, stromal cells_1, stromal cell_2, proliferating cells, stromal cells_3, pulmonary neuroendocrine cells (PNEC) and airway epithelial cells) for further analysis. We re-identified marker genes for the merged nine clusters and selected top positive marker genes per cluster for heatmap plot using the *DoHeatmap* function in the R Seurat package. The rest plots were generated using the R ggplot2 package.

### RNA-Seq before and following viral infections

Organoid infections were performed at an MOI of 0.1 and harvested at 24 hpi in DMEM supplemented with 0.3% BSA, 4.5 g/L D-glucose, 4 mM L-glutamine and 1 μg/ml TPCKtrypsin. Total RNA was extracted in TRIzol (Invitrogen) and DNase I treated using Directzol RNA Miniprep kit (Zymo Research) according to the manufacturer’s instructions. RNAseq libraries of polyadenylated RNA were prepared using the TruSeq RNA Library Prep Kit v2 (Illumina) or TruSeq Stranded mRNA Library Prep Kit (Illumina) according to the manufacturer’s instructions. cDNA libraries were sequenced using an Illumina NextSeq 500 platform. The resulting single end reads were checked for quality (FastQC v0.11.5) and processed using the processed using the nf-core RNA-seq (v.1.4.2) workflow. Samples had adapters trimmed using Trim Galore (v0.6.4) then ribosomal reads removed using SortMeRNA (v.2.1)before being aligned to human reference genome (GRCh38) using STAR aligner^30^ (v.2.6.1). Raw gene counts were quantified using Subread (v.1.6.4) featureCounts.

After further filtering and quality control, R package edgeR^31^ was used to calculate RPKM and Log2 counts per million (CPM) matrices as well as perform differential expression analysis. Principal component analysis was performed using Log2 CPM values and gene set analysis was run with WebGestalt ^32^. Heatmaps and bar plots were generated using Graphpad Prism software, version 7.0d.

### High Throughput Chemical Screening

hPSC-derived lung organoids were dissociated using TrypLE for 10 min in a 37°C waterbath and replated into 10% Matrigel-coated 384-well plates at 10,000 cells/40 μl medium/well. Six hour after plates, compounds from an in-house FDA-approved drug library (Prestwick) were added at 10 μM. DMSO treatment was used as a negative control. Two hours late, cells will be infected with SARS-CoV-2 pseudo virus (MOI=0.01). After 24 hpi, hPSC-COs were harvested for luciferase assay following the Luciferase Assay System protocol (Promega).

## QUANTIFICATION AND STATSTICAL ANALYSIS

N=3 independent biological replicates were used for all experiments unless otherwise indicated. n.s. indicates a non-significant difference. *P*-values were calculated by unpaired two-tailed Student’s t-test unless otherwise indicated. \**p*<0.05, \*\**p*<0.01 and \*\*\**p*<0.001.

## Acknowledgement

This work was supported by Department of Surgery, Weill Cornell Medicine (T.E., F.P, S.C.), and (NCI R01CA234614, NIAID 2R01AI107301 and NIDDK R01DK121072 and 1RO3DK117252), Department of Medicine, Weill Cornell Medicine (R.E.S.), by the Defense Advanced Research Projects Agency (DARPA-16-35-INTERCEPT-FP-006, B.T.) and by the Jack Ma Foundation (D.D.H). S.C and R.E.S. are supported as Irma Hirschl Trust Research Award Scholars. V.G. is a Weill Cornell Department of Medicine *Fund for the Future* awardee, supported by the Kellen Foundation. The authors would like to thank Dr. Harold Varmus at Weill Cornell Medicine for his support and Dr. Tom Moran, Center for Therapeutic Antibody Discovery at the Icahn School of Medicine at Mount Sinai for providing anti-SARS-CoV-SPIKE antibody.

## Data Availability

RNA-seq data is available from the GEO repository database with accession number GSE148697.

## Author Contribution

S. C., H. J. C., R.E.S., T. E., D. H., B. T., L.C., and H.W., conceived and designed the experiments. Y.H., L. Y., X. D., F. P., X.P., Z.Z., Y.B., J.L.J, D.N., F.D., and T.M.Y performed organoid differentiation, in vivo transplantation, pseudo-virus infection and drug screening.

P. W, Y. H., performed SARS2-CoV-2 pseudo-entry virus related experiments.

B. N., S.U., and B. T., performed SARS2-CoV-2 related experiments.

F.D., T. Z., J. X. Z., D. X., X. W., D.R., S.H., performed the scRNA-sequencing and bioinformatics analyses.

## Declaration of Interests

R.E.S. is on the scientific advisory board of Miromatrix Inc. The authors have no conflict of interest. L.C.C. is a founder and member of the board of directors of Agios Pharmaceuticals and is a founder and receives research support from Petra Pharmaceuticals. L.C.C. is an inventor on patents (pending) for Combination Therapy for PI3K-associated Disease or Disorder, and The Identification of Therapeutic Interventions to Improve Response to PI3K Inhibitors for Cancer Treatment. L.C.C. is a co-founder and shareholder in Faeth Therapeutics. T.M.Y. is a stockholder and on the board of directors of DESTROKE, Inc., an early-stage start-up developing mobile technology for automated clinical stroke detection.

**Table S3.** Antibodies used for immunocytochemistry, intracellular flow cytometry analysis and western blotting analysis.

| Usage | Antibody | Clone # | Host | Catalog # | Vendor | Dilution |
| --- | --- | --- | --- | --- | --- | --- |
| Immunocytochemistry | Human ACE-2 Antibody | Polyclonal | Goat | #AF933 | R&D Systems | 1:200 |
| Immunocytochemistry | Firefly luciferase Monoclonal Antibody (CS 17) | CS 17 | Mouse | #35-6700 | Thermo Fisher Scientific | 1:200 |
| Immunocytochemistry | Anti-NKX2.1 Antibody | Polyclonal | Rabbit | #WRA B-1231 | Seven Hills Bioreagents | 1:500 |
| Immunocytochemistry | Anti-SOX2 Antibody | Y-17 | Goat | #sc-17320 | Santa Cruz | 1:150 |
| Immunocytochemistry | Anti-FOXA2 Antibody | M-20 | Goat | #sc-6554 | Santa Cruz | 1:150 |
| Immunocytochemistry | Anti-SP-C Antibody | Polyclonal | Rabbit | #WRA B-76694 | Seven Hills Bioreagents | 1:500 |
| Immunocytochemistry | Anti-SP-B Antibody | Polyclonal | Rabbit | Cat# WRAB-48604 | Seven Hills Bioreagents | 1:500 |
| Immunocytochemistry | Anti-SARS-CoV-Spike antibody [2B3E5] |  | Mouse |  | Provided by Dr. Tom Moran | 1:100 |
| Immunocytochemistry | Donkey anti-Mouse IgG (H+L) Highly Cross-Adsorbed Secondary Antibody, Alexa Fluor 488 | Polyclonal | Donkey | #A-21202 | Thermo Fisher Scientific | 1:500 |
| Immunocytochemistry | Donkey anti-Rabbit IgG (H+L) | Polyclonal | Donkey | #A-21207 | Thermo Fisher Scientific | 1:500 |
|  | Secondary Antibody, Alexa Fluor 594 conjugate |  |  |  |  |  |
| Immunocytochemistry | Donkey anti-Goat IgG (H+L) Cross-Adsorbed Secondary Antibody, Alexa Fluor 647 | Polyclonal | Donkey | #A-21447 | Thermo Fisher Scientific | 1:500 |

**Table S3.**
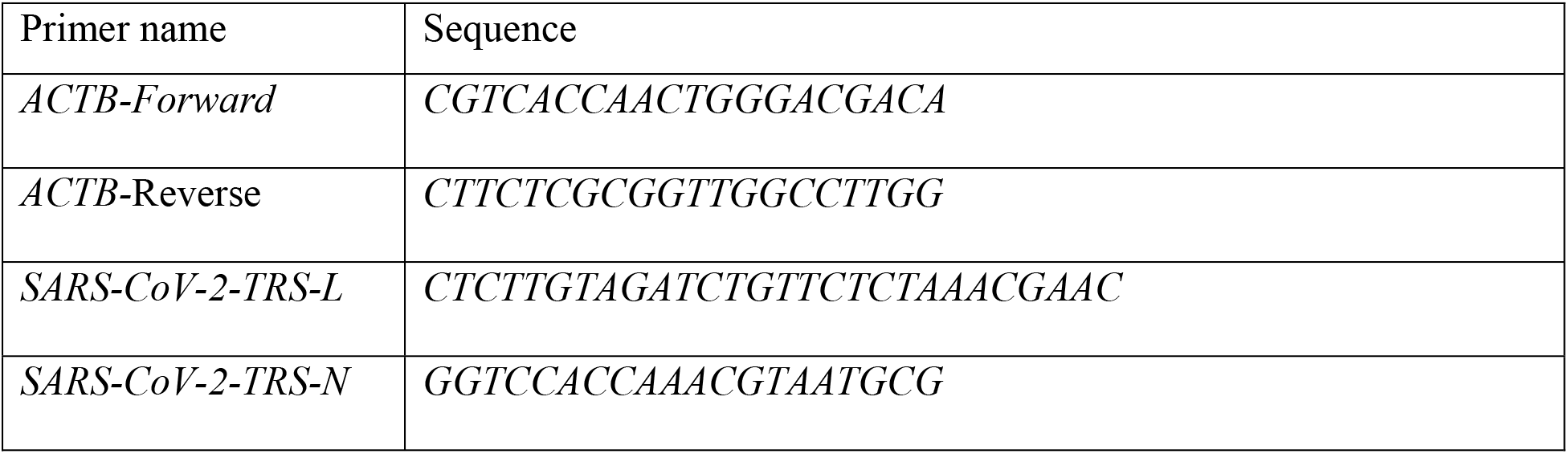
Primers used for qRT-PCR.

## Extended Data

**Extended Data Figure 1.**
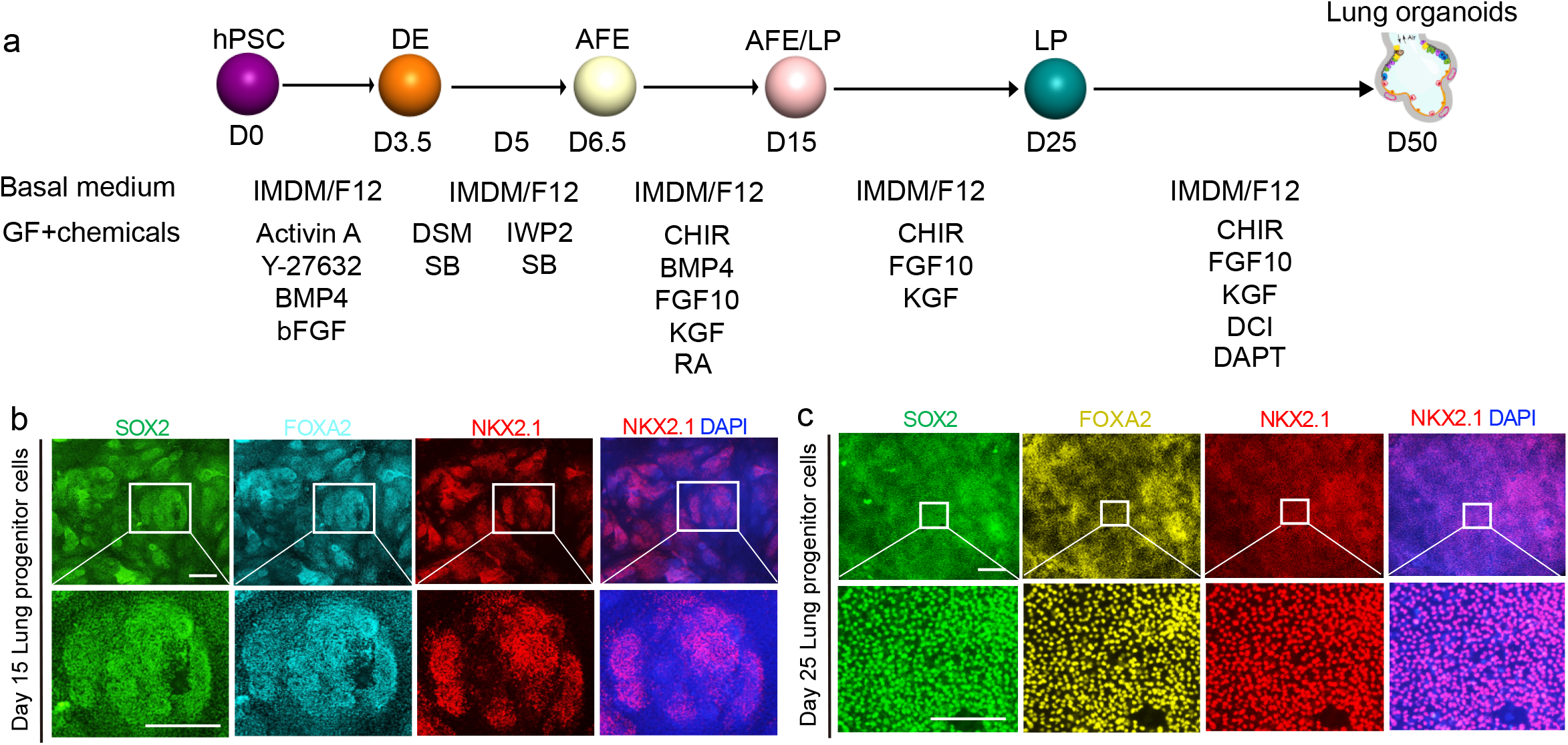
Directed differentiation of hPSC toward lung organoids. **a,** Scheme of directed differentiation of hPSCs to lung organoids. **b, c** Immunostaining was performed in the hPSC-derived cell cultures at day 15 (b) and day 25 (c). Scale bars=100 μm.

**Extended Data Figure 2.**
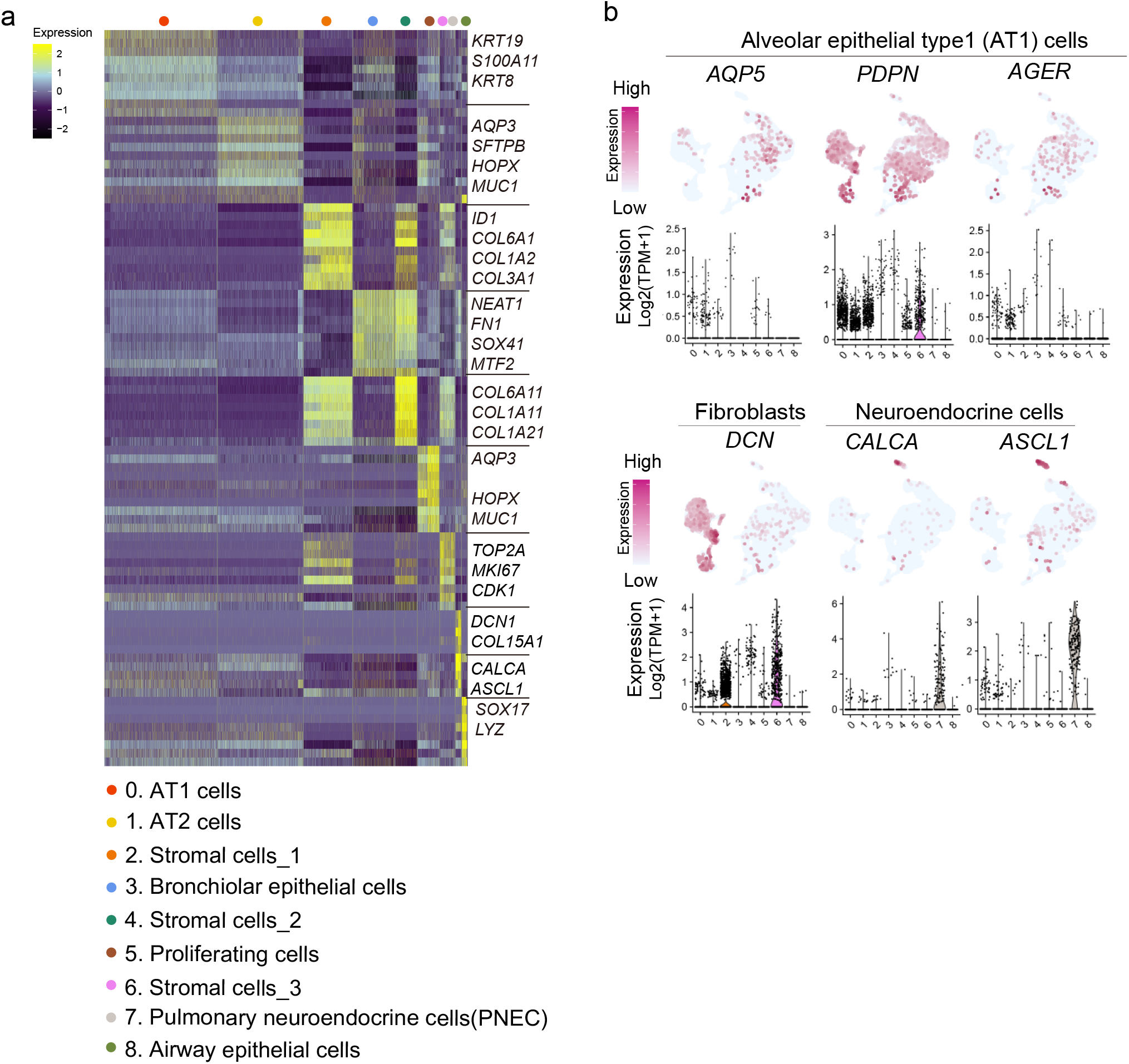
Single cell RNA-seq analysis of hPSC-derived lung organoids. **a,** Heatmap of enriched genes in each cluster of scRNA profiles in hPSC-derived lung organoids. Each row represents one top differentially expressed gene and each column represents a single cell. **b.** Putative AT1, fibroblast and PNECs markers in each cluster in UMAPs. Relative expression of each marker gene range from low (light blue) to high (pink) as indicated. Individual cells positive for lung cell markers are donated by red dots. The violin plot shows the expression level (log2(TPM+1)) of indicated gene in each cluster.

**Extended Data Figure 3.**
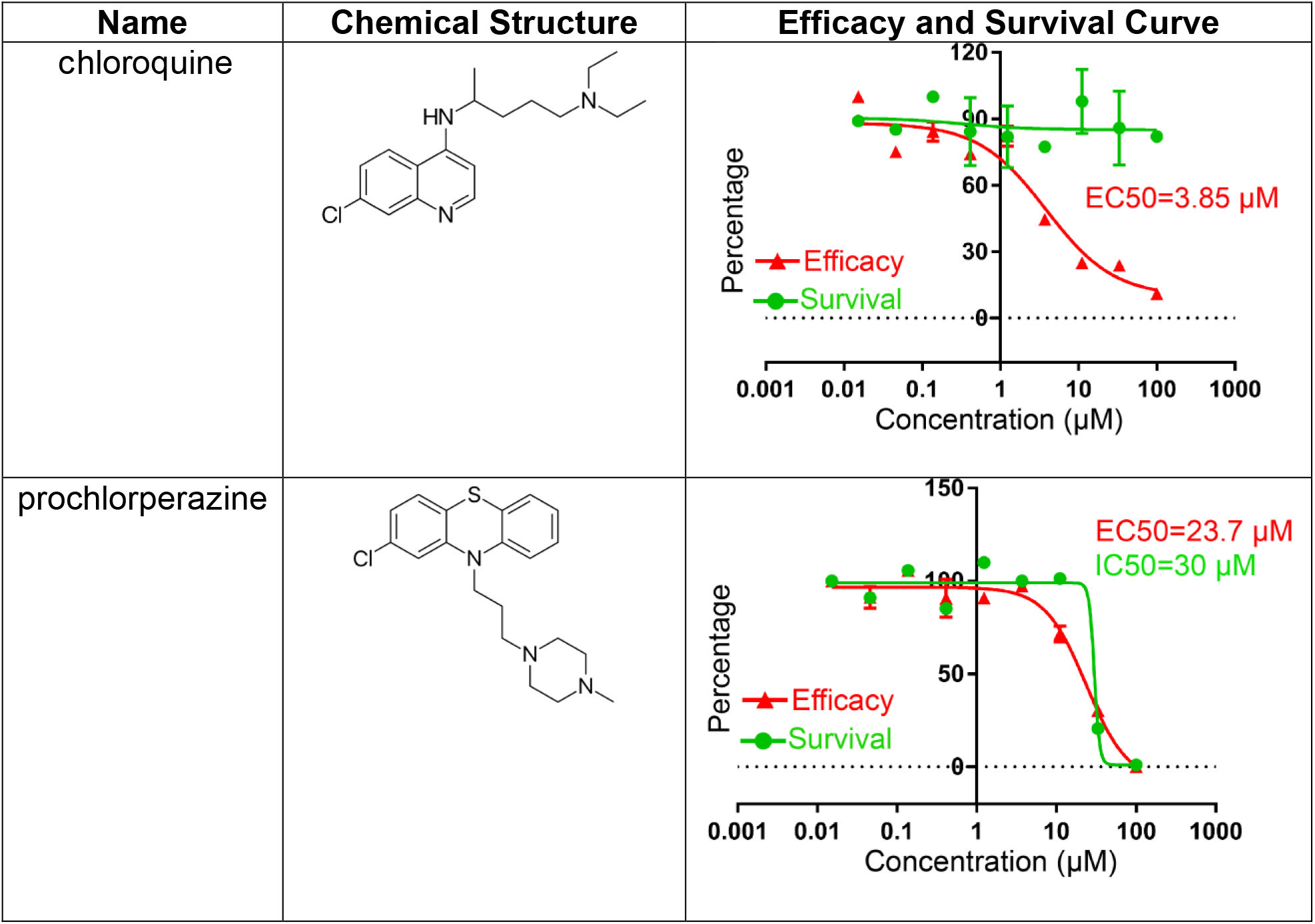
Chemical structure, efficacy curve and toxicity curve of primary hit drug candidates.

**Extended Data Figure 4.**
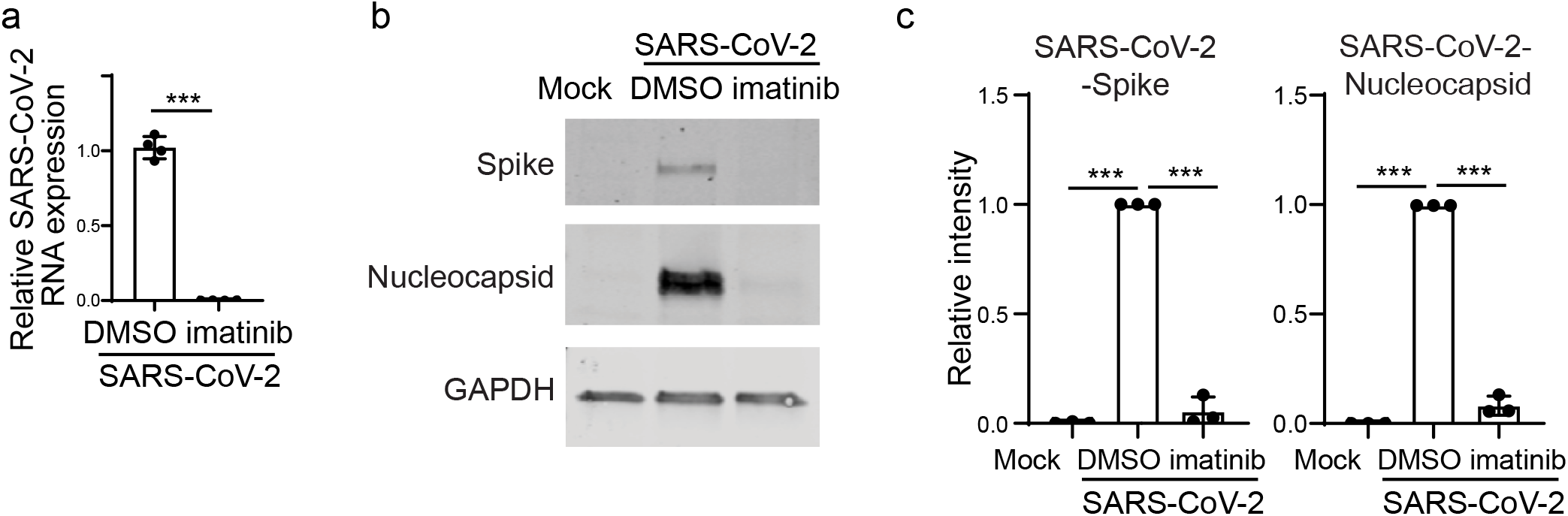
Imatinib shows anti-SARS-CoV-2 activity on Vero cells. **a,** qRT-PCR analysis of DMSO or 10 μM imatinib treated Vero cells at 24 hpi (SARS-CoV-2, MOI=0.01). Data was presented as mean ± STDEV. *P* values were calculated by unpaired two-tailed Student’s t test. \**P* < 0.05, \*\**P* < 0.01, and \*\*\**P* < 0.001. **b, c,**Western blotting (b) and quantification (c) of DMSO or 10 μM imatinib treated Vero cells at 24 hpi (SARS-CoV-2, MOI=0.01).

